# β-glucan imprinting remodels macrophage function in response to environmental cues

**DOI:** 10.1101/2021.08.30.458241

**Authors:** Alícia C. Piffer, Giorgio Camilli, Mathieu Bohm, Rachel Lavenir, Jessica Quintin

## Abstract

*In vitro*, exposure of human primary monocytes to the fungal β-glucan enhances their pro-inflammatory responsiveness towards several pathogens. Yet, the role of environmental condition of this process remains unclear. Here we found that β-glucan-induced innate immune memory counteract the anti-inflammatory status of the macrophages. In response to β-glucan imprinting, M-CSF-(M2-like-) macrophages increase their pro-inflammatory responsiveness to secondary stimuli associated with decrease of the M-CSF differentiation hallmarks. In contrast, in GM-CSF-(M1-like-) environment, β-glucan imprinting reduced the pro-inflammatory canonical feature of the macrophages, together with their terminal differentiation marks. Comparing M-CSF and GM-CSF environment, we observed that β-glucan-imprinted macrophages present comparable functions in terms of cytokine responses, phagocytosis, oxidative burst, and angiogenesis. This effect is mediated through Dectin-1 and associated with altered expression of the master regulators of macrophage terminal differentiation, IRF5 and IRF3. β-glucan-induced innate immune memory skews the commitment of the macrophages in complex environment towards similar and less terminally differentiated cells. Together, these observations suggest a potential therapeutic role for β-glucan-induced modulation of innate memory in different pathological contexts.

## Introduction

Immunological memory as an exclusive hallmark of adaptive immune response is now outdated. Numerous reports demonstrate that innate immune cells can recall a previous encounter with pathogens or their compounds, such as the fungal β-1,3-glucan, and generate innate immune memory (1–3). Over the past few years, we and others have shown that β-glucan imprinting of human primary monocytes cultured in human serum enhances their pro-inflammatory responsiveness towards several other pathogens that are not necessarily fungal-related (1, 4, 5). This imprinted innate memory associated with enhanced pro-inflammatory state has been termed “trained immunity” (6, 7).

While these studies on monocytes have been crucial to demonstrate innate immune memory and identify underlying mechanisms, bloodstream classical monocytes are not themselves effector cells during infection, but rather mostly circulating intermediates. Indeed, circulating monocytes infiltrate peripheral tissues, where they are recruited and contribute to effective control and clearance of viral, bacterial, fungal, or protozoal infections. In tissues, following conditioning by local growth factors, pro-inflammatory cytokines, and microbial products, they differentiate into macrophages. Depending on the microenvironmental stimuli, macrophages undergo activation or “polarisation” along a wide spectrum of phenotypes (8). Hence the *in vivo* interaction of β-glucan with the innate cells would occur in a complex environment.

Recently, by recapitulating different physiological environments *in vitro* we uncovered a novel property of innate immune memory, in which β-glucan imprints on differentiating macrophages an inhibitory effect on NLRP3 inflammasome activation and subsequent mature IL-1β secretion that is independent of previously described transcriptional regulation mechanisms (9). This underscores that we do not yet fully comprehend the effects of β-glucan imprinting on polarising and differentiating macrophages in complex environment. Understanding the potential of β-glucan to induce innate immune memory in a complex physiological environment is an important first step towards harnessing the beneficial effects of innate immune memory in future translational research.

Granulocyte-macrophage colony-stimulating factor (GM-CSF)-differentiated human macrophages are polarised to classically activated macrophages, also known as M1-like macrophages, which harbour immune-stimulatory properties against infections with high expression of pro-inflammatory cytokines, high ROS production, and cytotoxic function against tumour cells (10). M-CSF-differentiated human macrophages are polarised to alternatively activated, or M2-like, macrophages, described as an anti-inflammatory cell type, with low cytotoxic function therefore potent pro-tumoral activity but with high tissue-remodelling and repair activity (11).

In this study, we investigated the specific effects of β-glucan imprinting on polarising human macrophages. We show that not only does β-glucan-induced innate immune memory enhance responsiveness and function of macrophages but under certain conditions can reduce it. Responses to secondary challenges were increased by β-glucan imprinting in M-CSF-polarising macrophages but decreased in GM-CSF-polarising cells. Macrophages were less differentiated upon β-glucan imprinting as the hallmarks of terminally matured macrophages were all distinctively decreased. Of interest, M-CSF- and GM-CSF- β-glucan-imprinted macrophages present similar cytokine responses to secondary infection, phagocytosis, angiogenesis and ROS signalling functions. The β-glucan imprinting effect on the polarisation of the macrophages is mediating by the Dectin-1 receptor and modify the expression of the master regulators IRF5 and IRF3. Finally, β-glucan-induced innate memory skew the terminal differentiation of macrophages early in the differentiation process. Altogether, these observations provide new insights into the mechanisms of β-glucan-induced innate memory with therapeutics implications.

## Materials and Methods

### Cell isolation

Human PBMCs were separated on Ficoll-Paque (GE Healthcare) by density centrifugation of heparinized blood from healthy donors (“Etablissement Français du Sang” EFS, Paris, France). Monocytes were purified by positive selection using the CD14 MicroBeads, human (Miltenyi Biotec). The purity evaluated by flow cytometry was routinely >90%.

### Macrophage differentiation

Monocytes were seeded at 10^6^ cells/mL in RPMI 1640, supplemented with 2 mM GlutaMax, 50 μg/mL gentamicin and 1 mM sodium pyruvate (Gibco) and incubated 24h with GM-CSF (25 ng/mL, 130-093-865, Miltenyi Biotec) or M-CSF (25 ng/mL, 216-MC-025, R&D Systems) in the presence or absence of purified β-glucan from *Candida albicans* (10 μg/mL unless otherwise stated; depyrogenated and kindly provided by Dr. David L. Williams, East Tennessee State University). After the first 24 hours, the cells were washed with warm PBS and incubated for 5 more days in RPMI containing 10% FBS (F7524, Sigma-Aldrich) in the presence of GM-CSF or M-CSF. The medium was changed once at day 3. Cells were maintained in a humidified atmosphere of 5% CO_2_ at 37°C.

For Dectin-1 inhibition, anti-Dectin-1 mAb (10 μg/mL, ab82888, abcam) or IgG1 Isotype control (10 μg/mL, 401402, Biolegend) was added 30 min prior to β-glucan stimulation and maintained in culture medium for 24h. After this period, cells were differentiated as described above.

For evaluation of M-CSF effect on β-glucan imprinting, monocytes were seeded at 10^6^ cells/mL in RPMI 1640, supplemented as described previously and incubated 24h with GM-CSF (25 ng/mL) in the presence of M-CSF (1000 pg/mL, 100 pg/mL or 50 pg/mL), or β-glucan (10 μg/mL) or medium as control. Then, the cells were washed with warm PBS and incubated for 5 more days in RPMI containing 10% FBS in the presence of GM-CSF. The medium was changed once and the cells were maintained in a humidified atmosphere of 5% CO_2_ at 37°C.

### Viability assay

Monocytes were seeded in black 96-well plates and differentiated in macrophages as described above. At day 6, one set of macrophages was treated with saponin 0.05% for 15 minutes (positive control for PI). Then, the cells were washed three times with warm PBS and incubated with fluorescein diacetate (8 μg/mL, Sigma-Aldrich) and propidium iodide (20 μg/mL, Sigma-Aldrich) diluted in RPMI 1640 for 5 minutes in the dark. Then cells were washed three times with warm PBS and were imaged using an EVOS FL system (Thermo Fisher Scientific, Ex: 470nm, Em: 525nm for GFP and Ex: 530nm, Em: 592nm for PI) and PI fluorescence was measured with a Synergy H1 microplate reader (Ex: 520nm, Em: 610nm; BioTek Instruments).

### Cytokine assay

Macrophages were differentiated as described above and, at day 6 (unless otherwise stated), were stimulated with heat-inactivated *Staphylococcus aureus* (5×10^7^ cells/mL) or heat-inactivated *Escherichia coli* (1×10^5^ cells/mL) for 24 hours in RPMI. IL-6, TNF-α, IL-12p40 (R&D Systems) and IL-10 (BioLegend) levels were determined by enzyme-linked immunosorbent assays (ELISA) on cell culture supernatants, according to the manufacturer’s instructions. Absorbance was measured with a Synergy H1 microplate reader (BioTek Instruments).

### Reactive oxygen species measurement

Monocytes were seeded in white 96-well plates and differentiated as described above. At day 6, macrophages were stimulated with 500 ng/mL of PMA (phorbol 12-myristate 13-acetate, Sigma-Aldrich) diluted in HBSS (Hank’s Balanced Salt Solution – Gibco). At the same time, Luminol (1mM; Sigma-Aldrich), also diluted in HBSS, was added, and light emission was measured every 2 minutes for a period of 120 min using the Synergy H1 microplate reader (Biotek Instruments, USA). The light emission levels are expressed as RLU (Relative Light Units = luminescence measured by the luminometer).

### Phagocytosis assay

To evaluate the phagocytic activity of macrophages, pHrodo Green inactivated *S. aureus* BioParticles conjugate was used (P35367, Life Technologies). Monocytes were seeded in a flat clear bottom black 96-well plate (3603, Corning) and differentiated as described above. On the sixth day of macrophage differentiation, pHrodo BioParticles were resuspended in Live Imaging Solution (Life Technologies), sonicated and incubated with macrophages (5 μg/mL final concentration). The plate was centrifuged 300g for 2 minutes to synchronise the interaction between the BioParticles and macrophages and incubated for 1 hour at 37 °C. After the incubation time, fluorescence was measured with a Synergy H1 microplate reader (Ex: 505 nm, Em: 535 nm; BioTek. Instruments, USA). The fluorescence emission levels are expressed as RFU (Relative Fluorescence Units).

Phagocytosis was also measured by flow cytometry: monocytes were seeded in a 12-well plate and differentiated as described above. On the sixth day of macrophage differentiation, the macrophages were stimulated with pHrodo BioParticles as described above, for 30 minutes at 37°C. Then cells were detached from the plate using Accutase solution (Sigma-Aldrich) at 37°C for 15 minutes followed by manual shaking and repeated pipetting. Acquisition was performed on a LSR Fortessa (BD Biosciences) and *S. aureus* BioParticles were detected in the FITC channel. Analysis was performed using FlowJo Software (Treestar, Ashland). Phagocytosis rate was quantified as the percentage of cells positive for the *S. aureus* BioParticles (%FITC+) and phagocytosis efficacy, or the average amount of bacteria per cell, was quantified as the Mean Fluorescence Intensity (MFI) for *S. aureus* BioParticles within FITC-positive cells.

### RNA isolation and real-time PCR analysis

Macrophages were differentiated as described above and, at day 6 were stimulated for 16 hours with heat-inactivated *S. aureus* (5×10^7^ cells/mL) or for 3 hours with heat-inactivated *E. coli* (1×10^5^cells/ml) in RPMI, and total RNA was isolated from macrophages using TRIzol reagent (Invitrogen). RNA quality was monitored by spectrophotometric analysis. Total RNA (1 μg) from each sample was used for reverse transcription with the High-Capacity cDNA Reverse Transcription Kit (Applied Biosystems, Life Technologies). Real-time PCR was performed in ABI Step One Plus Real-Time PCR System (Applied Biosystems, Life Technologies) using the TaqMan 2× Universal Master Mix and the following TaqMan Gene Expression Assays: IL-12, Hs01011518, CCL22, Hs01574247; CCL2, Hs00234140; CXCL9, Hs00171065; VEGFA, Hs00900055; IL-6, Hs00174131 (Applied Biosystems, Life Technologies). Each sample was assayed in triplicates.

Thermal cycling conditions were as follows: denaturation at 95°C for 10 min and 40 cycles of 95°C for 15 sec and 60°C for 60 sec. Fold change of the cytokine genes in the samples was calculated using the 2^−ΔΔCT^ method. All values were normalised to endogenous control β-Actin (Hs99999903; Applied Biosystems, Life Technologies) and were expressed in arbitrary units.

### Flow cytometry

Monocytes were seeded in a 12-well plate and differentiated as described above. At day 6, macrophages were detached from the plate using Accutase solution (Sigma-Aldrich) at 37°C for 15min followed by manual shaking and repeated pipetting. Fc receptors were blocked by incubation with FcR Blocking Reagent (Miltenyi Biotech) and staining was performed using antibodies purchased at Miltenyi Biotech: anti-CD163 PE (clone REA812, Cat# 130-112-128,), anti-CD206 PE-Vio770 (clone DCN228, Cat# 130-100-152) and anti-CD14 VioGreen (clone REA599, Cat# 130-110-524). Cells were also stained with Fluorescence Minus One (FMO) mixes containing the relevant isotype control: REA control, clone REA293, coupled with VioGreen (Cat# 130-113-442), or PE (Cat#130-113-438), or mouse IgG1 PE-Vio770 (clone IS5-21F5, Cat# 130-113-202). Acquisition was performed with a LSR Fortessa (BD Biosciences) after fixation of cells with 4% Formaldehyde Solution at room temperature protected from light. Analysis was performed using FlowJo Software (Treestar, Ashland) with the following gating strategy: cells were separated from debris using Forward (size) and Side Scatter (granularity) channels, and doublets were excluded using Forward Scatter Area versus Width. Mean Fluorescence Intensity measures were taken from these selected single cells.

### Transmigration assay

Monocytes were seeded in a 24-well plate and differentiated as described above. On the sixth day of macrophage differentiation, cells were washed once with warm PBS and stimulated, for 24 hours, with heat-inactivated *Staphylococcus aureus* (5×10^7^ cells/mL) resuspended in Endothelial medium (C-22210, PromoCell) complemented with 10% of fetal bovine serum. After stimulation time, the conditioned medium was placed in the bottom chamber of a transwell system (pore size 5 μm, Costar). HUVECs cells (2.5×10^5^ cells/mL - PromoCell), resuspended in free endothelial medium, were placed in the top chamber and the system was incubated for 24 hours in a humidified atmosphere of 5% CO_2_ at 37°C. Then, the chamber was washed twice with warm PBS, the cells were fixed with 4% Formaldehyde Solution at 37 °C for 15 minutes. Then, the cells were permeabilized with 0,5% Triton X-100 for 10 minutes at room temperature and stained 10 minutes in the dark with DAPI (Thermo Fisher Scientific). To only count the cells that migrated through the pores, the cells in the top chamber were eliminated with a swab. The transwell chambers were imaged using EVOS FL system (Ex: 360nm, Em: 447nm for DAPI, Thermo Fisher Scientific) and migrated cells were counted using ImageJ software (NIH, Bethesda, MD, USA).

### Western blot

Monocytes were differentiated as described above and, after 6 days of differentiation, cells were stimulated with heat-inactivated *Staphylococcus aureus* (5×10^7^ cells/mL) for 24 hours. After stimulation, total cellular protein extracts were obtained by lysing cells for 30 min at 4°C in lysis buffer (Pierce RIPA Buffer) in the presence of protease inhibitors (cOmplete Protease Inhibitor Cocktail, ROCHE). Samples were centrifuged for 20 min at 4°C, 13,000 rpm and supernatants were collected in fresh tubes. Proteins from cell extracts were resuspended in Laemmli Buffer 1X, boiled at 95°C for 7 min and resolved by SDS-PAGE. Then, proteins were transferred with an iBlot instrument (Life Technologies) to a nitrocellulose membrane. Immunoblotting was performed using the following primary antibodies: anti-IRF3 (sc-33641, Santa Cruz Biotechnology), anti-GAPDH (sc-25778, Santa Cruz Biotechnology) and anti-IRF5 (20261S, Cell Signaling). Anti-mouse or anti-rabbit IgG, HRP-linked antibodies (NA931V and NA934V respectively, GE Healthcare) were used as secondary antibodies, and protein bands were visualized with ECL Prime (GE Healthcare) or SuperSignal West Femto Maximum Sensitivity Substrate (Thermo Fisher Scientific) in a myECL imager (Thermo Fisher Scientific). Proteins were normalised for GAPDH using Thermo Scientific myImageAnalysis Software.

### Statistic

The results were analysed using a Wilcoxon matched-pairs signed-rank test (unless otherwise stated). A *p* value below 0.05 was considered statistically significant. All data were analysed using GraphPad Prism software. Data are shown as mean ± standard error of the mean.

## Results

### β-glucan-induced innate immune memory alter the anti-inflammatory and differentiation hallmarks of M-CSF macrophages

Previous works that have studied the innate immune memory (1, 4, 12, 13), and more specifically the enhanced secretion of IL-6 and TNFα upon secondary stimulation, a hallmark of trained immunity, have used isolated human monocytes from PBMCs and maintained them in human serum only. Human serum contains low but detectable amounts of M-CSF (around 2 to 10 ng/mL (14)) and human monocytes isolated from PBMCs can differentiate into macrophages with as little as 10 ng/mL of M-CSF. It is therefore likely that macrophages obtained in human serum only are comparable to M-CSF-macrophages (**Figure 1A**).

**Figure 1.**
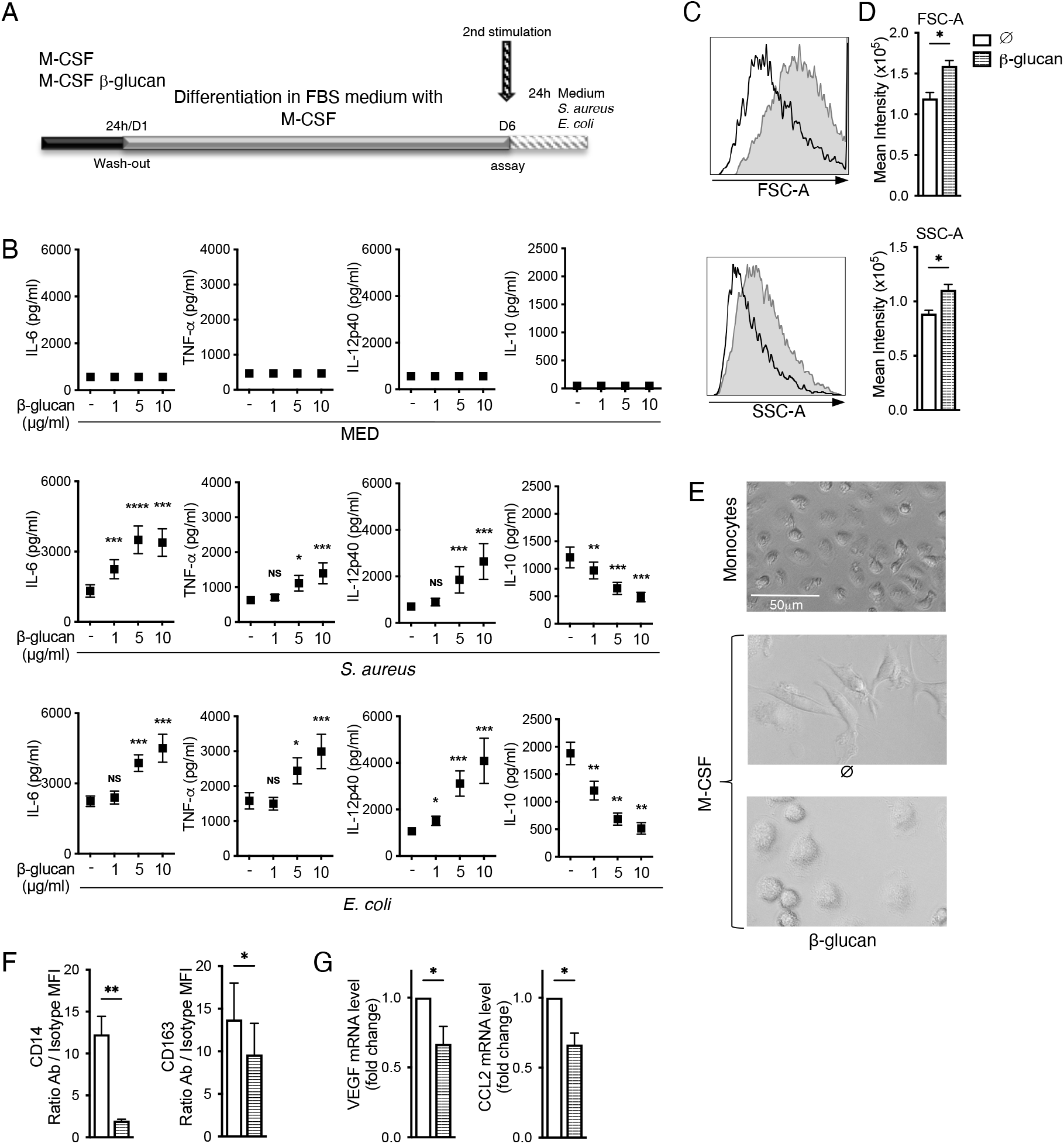
β-glucan imprinting increases the pro-inflammatory status of M-CSF macrophages and reduce their terminal differentiation marks. (A) Schematic overview of the methodology used in this study. Monocytes were preincubated with β-glucan or left untreated in a medium containing M-CSF. After 24 hours, the stimulus was washed away, and the cells were differentiated for an additional 5 days, after which the macrophages were re-stimulated. (B) Macrophages obtained in M-CSF environment were re-stimulated with heat-inactivated *Staphylococcus aureus* or *Escherichia coli*. Culture supernatants were collected after 24h and concentrations of secreted TNF-α, IL-6, IL-12, and IL-10 were determined by ELISA. (C, D) Monocytes at day 0 and M-CSF macrophages at day 6 were detached and analysed for size (FSC-A) and granularity (SSC-A) by flow cytometry. (C) Representative flow cytometry histograms showing FSC-A (top) and FSC (bottom) of macrophages. White filled histograms: naïve macrophages, grey filled histograms: β-glucan-treated macrophages. (D) Histogram of the size (FSC-A, top) and granularity (SSC-A, bottom) mean intensity quantification. (E) Monocytes at day 0 and M-CSF macrophages at day 6 were also imaged to evaluate their morphology. (F) At day 6 of the differentiation process, macrophages were stained for the indicated cell surface markers, or with FMO antibody mixes containing the relevant isotype controls, for flow cytometry analysis. Intensity of expression of a given marker was measured by doing the ratio of the full staining MFI over the isotype MFI (‘ratio MFI’). (G) At day 6 of differentiation, macrophages were also re-stimulated for 18h with heat-inactivated *S. aureus* and *VEGF* and *CCL2* gene expression was analysed by Real-Time quantitative PCR (RT-qPCR). Results were normalised to β-actin expression levels. The results obtained for the cells differentiated with M-CSF in the absence of β-glucan were arbitrarily set at 1 to express the results as fold change. Graphs show the mean ± SEM of at least three independent experiments. For B *n* = 11; C, D, *n*=6; F, *n* = 6-8; G, *n* = 7; \**p* < 0.05, \*\**p* < 0.01, \*\*\**p*<0.001. Wilcoxon matched-pairs signed-rank test (*: cells stimulated with β-glucan vs. non stimulated with β-glucan).

As expected for M-CSF-derived macrophages (9), β-glucan imprinting increased their pro-inflammatory status and responses to gram-positive and gram-negative bacteria challenges in a dose-dependent manner (**Figure 1B**). These effects were also observed at the transcriptional level (**Supplementary Figure 1A**) as previously reported for β-glucan-imprinting of monocytes. Although past literature demonstrated that β-glucan inhibits spontaneous human monocyte apoptosis in minimum medium (5) explaining the decreased cell number observed in minimum medium (4), we have previously established that β-glucan together with growth factors does not alter the viability of macrophages (9). In addition, using the strongest effective concentration of β-glucan for imprinting (10 μg/mL), we further ascertained that β-glucan does not increase survival of M-CSF-macrophages in this study (**Supplementary Figure 2A, B**). In minimum medium, β-glucan-treated-monocytes were reported to not only change their cytokines response to secondary stimuli but also their morphology (4). Specifically, when monocytes were incubated for 24 h and rested for 6 days in human-serum only, β-glucan-induced cells were bigger than their naive counter parts. We observed similar morphological changes in rich environment in which β-glucan-M-CSF-macrophages were bigger than their naïve counterpart but also more granular (**Figure 1C, D**). Of interest, cell morphology is one of the described polarisation features distinguishing environment-specific-macrophages (15) and upon β-glucan imprinting, M-CSF-macrophages did not reach their canonical spindle shape (**Figure 1E**) suggesting an alteration in their differentiation.

We therefore look at differentiation markers of M-CSF-macrophages. With respect to cell surface markers of macrophages, the LPS coreceptor CD14 is a monocyte-specific surface marker whose expression ultimately decreases upon differentiation into macrophages but remains more expressed in M-CSF-macrophages compared to GM-CSF-macrophages (16). β-glucan-imprinted M-CSF-macrophages presented a lower CD14 surface expression, compared to their non-primed counterparts (**Figure 1F**). Similarly, β-glucan imprinting decreased the M-CSF-associated cell surface marker CD163 expression (**Figure 1F**). Finally, the M-CSF-associated markers *VEGF* and *CCL2* (17) expression decreased in M-CSF-macrophages after β-glucan imprinting (**Figure 1G**). Overall, it appears that in M-CSF anti-inflammatory environment, β-glucan imprinted increase the pro-inflammatory status of the macrophages to secondary infections, which is also associated with decreased hallmarks of M-CSF terminal differentiation.

### β-glucan-induced innate immune memory decrease the pro-inflammatory features of GM-CSF macrophages towards less differentiated macrophages

In order to understand the effect of β-glucan in diverse complex environment we repeated our investigation using an opposite environment to M-CSF, namely GM-CSF (**Figure 2A)** (10, 11, 18). Whereas the increase pro-inflammatory responses in imprinted-M-CSF-derived macrophages was detected as expected, we however interestingly observed that β-glucan imprinting of GM-CSF-differentiating macrophages decreased their responsiveness, resulting in less inflammatory macrophages (**Figure 2B**). Their pro-inflammatory cytokine responses to both gram-positive and gram-negative bacteria challenges decreased in a dose-dependent manner (**Figure 2B**). GM-CSF macrophages do not produce the anti-inflammatory cytokine IL-10 upon stimulation and no imprinting effects were observable. These regulations of pro-inflammatory cytokines secretion also affected the genes expression (**Supplementary Figure 1B**) excluding a post-translational regulation by β-glucan-reprogramming as we recently reported for the inflammasome-dependent IL-1β (9). In addition, the decrease responsiveness was not due to a decrease viability of the GM-CSF-macrophages (**Supplementary Figure 2A, B**). Monocytes stimulated by β-glucan do not release GM-CSF but secrete M-CSF (from 20 pg/mL up to 100 pg/mL depending on the amount of β-glucan) (5). Hence, the effect of β-glucan on GM-CSF-differentiating macrophages could be due to the imbalance in growth factors concentration. However, neither physiologically secreted nor high doses of exogenous M-CSF affected the cytokine responses of GM-CSF-macrophages to secondary bacterial stimulation (**Supplementary Figure 2C**). Therefore, the decreasing effect of β-glucan-induced innate immune memory on the pro-inflammatory status of GM-CSF differentiating macrophages was neither due to an effect of β-glucan on the viability of the macrophages nor to an imbalanced M-CSF concentration possibly caused by β-glucan stimulation.

**Figure 2.**
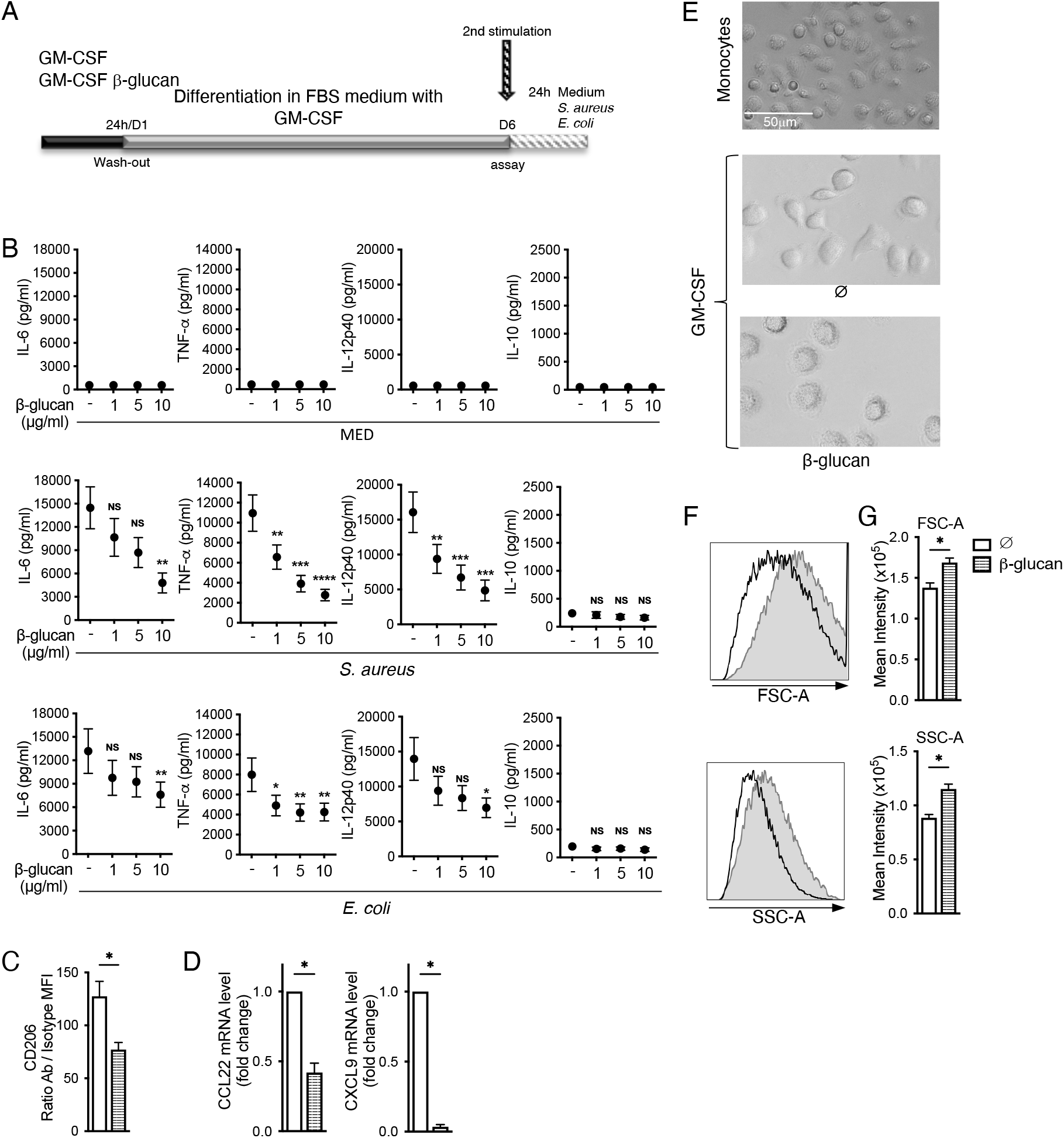
β-glucan imprinting decreases the pro-inflammatory and differentiation hallmarks of macrophages in GM-CSF environment. (A) Schematic overview of the methodology used in this study. Monocytes were preincubated with β-glucan or left untreated in a medium containing GM-CSF. After 24 hours, the stimulus was washed away, and the cells were differentiated for an additional 5 days, after which the macrophages were re-stimulated. (B) Macrophages were re-stimulated with heat-inactivated *Staphylococcus aureus* or *Escherichia coli*. Culture supernatants were collected after 24h and concentrations of secreted TNF-α, IL-6, IL-12, and IL-10 were determined by ELISA. (C) At day 6 of the differentiation process, macrophages were stained for the CD206 cell surface marker, or with FMO antibody mix containing the relevant isotype control, for flow cytometry analysis. Intensity of expression was measured by doing the ratio of the full staining MFI over the isotype MFI (‘ratio MFI’). (D) At day 6 of differentiation, macrophages were also re-stimulated for 18h with heat-inactivated *S. aureus* and *CCL22*, and *CXCL9* gene expression was analysed by Real-Time quantitative PCR (RT-qPCR). Results were normalised to β-actin expression levels. The results obtained for the cells differentiated with GM-CSF in the absence of β-glucan were arbitrarily set at 1 to express the results as fold change. (E) Monocytes at day 0 and GM-CSF macrophages at day 6 were imaged to evaluate their morphology. (F, G) Monocytes at day 0 and GM-CSF macrophages at day 6 were also detached and analysed for size (FSC-A) and granularity (SSC-A) by flow cytometry. (F) Representative flow cytometry histograms showing FSC-A (top) and FSC (bottom) of macrophages. White filled histograms: naïve macrophages, grey filled histograms: β-glucan-treated macrophages. (G) Histogram of the size (FSC-A, top) and granularity (SSC-A, bottom) mean intensity quantification. Graphs show the mean ± SEM of at least three independent experiments. For B *n* = 11; C, *n* = 6-8; D, *n* = 7; F,G, *n*=6 \**p* < 0.05, \*\**p* < 0.01, \*\*\**p*<0.001. Wilcoxon matched-pairs signed-rank test (*: cells stimulated with β-glucan vs. non stimulated with β-glucan).

The decrease in the canonical pro-inflammatory responses of GM-CSF macrophages upon β-glucan imprinting was also associated with decrease in their specific markers defining GM-CSF-macrophages terminal differentiation. As such, the GM-CSF-associated cell surface marker CD206 expression (**Figure 2C**) (19) and *CCL22* and *CXCL9* GM-CSF-markers decreased upon β-glucan-reprogramming in GM-CSF-macrophages (**Figure 2D**). Of note, the shape of the GM-CSF-macrophages was also altered (**Figure 2E**) with an increase in both size and granularity in β-glucan-imprinted macrophages compared to their non-treated counterpart (**Figure 2 F, G**).

Overall, it appears that β-glucan reprogramming specifically affect macrophages depending on the environment and do not always increase their responsiveness. In pro-inflammatory polarising environment, β-glucan-induced innate memory decreases their inflammatory status towards less differentiated macrophages.

### β-glucan reprogramming differentially affects macrophages functions depending on the environment towards common type of macrophages

We observe that β-glucan reprogramming differentially imprints pro-inflammatory GM-CSF- and anti-inflammatory M-CSF-macrophages. Comparing the β-glucan-imprinted macrophages obtained after a week, it appears that the opposite effects depending on the environment lead toward a similar intermediate type of macrophage that presents both pro- and anti-inflammatory responses to secondary infections (**Figure 3A**), suggesting an increase plasticity of the cells in terms of the type of response required.

**Figure 3.**
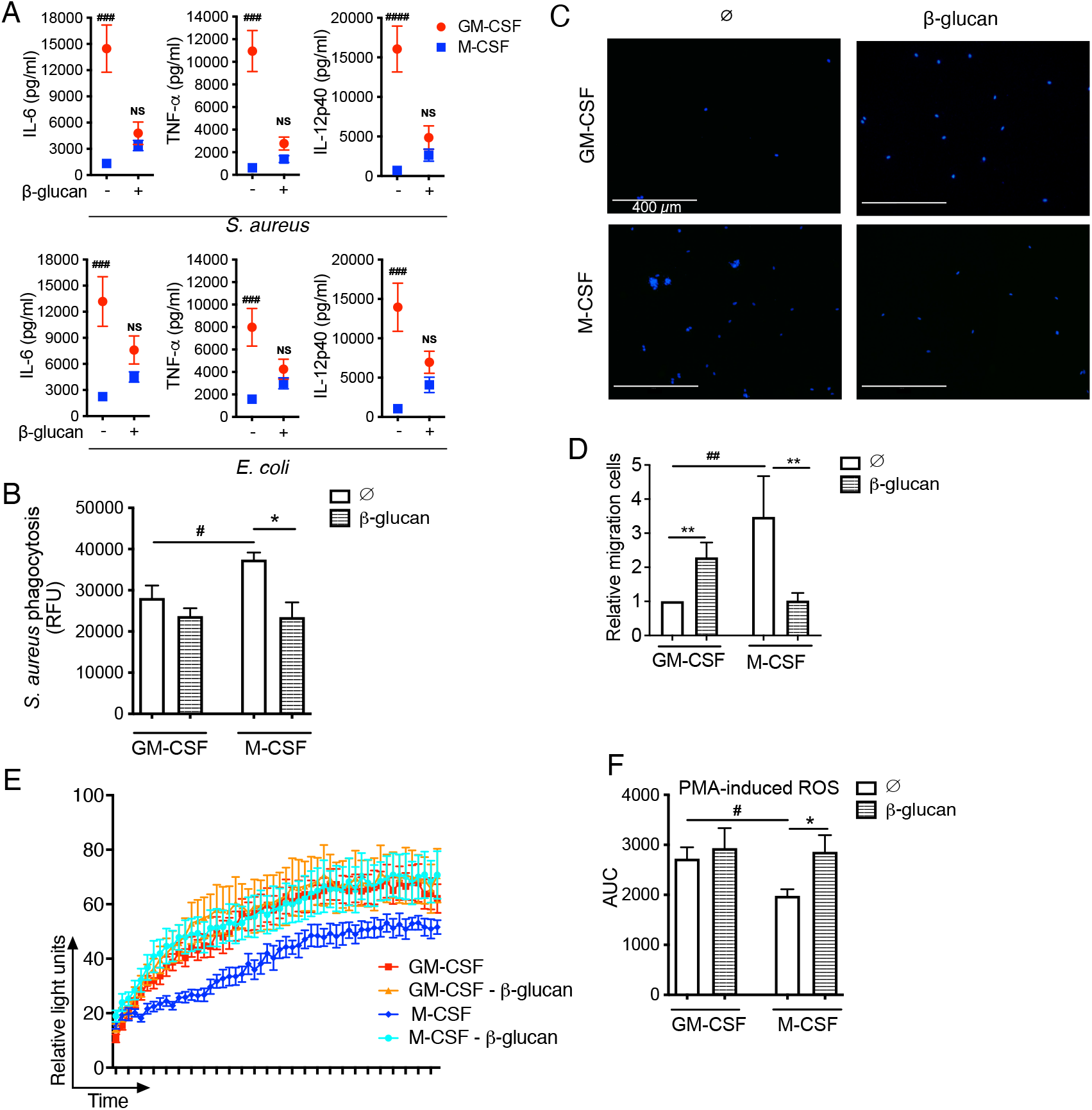
β-glucan imprinting oppositely affect macrophages depending on the environment leading towards macrophages with affects with similar functional activities. Monocytes were preincubated in the presence or absence of β-glucan and then differentiated in macrophages with either GM-CSF or M-CSF. (A) Macrophages were re-stimulated with heat-inactivated *Staphylococcus aureus* or *Escherichia coli*. Culture supernatants were collected after 24h and concentrations of secreted TNF-α, IL-6, and IL-12, were determined by ELISA. (B) After 6 days of culture, resulting macrophages were re-stimulated with pHrodo™ Green *S. aureus* Bioparticles™ for 1 hour. After the incubation time, phagocytosis was measured by reading the fluorescence and is expressed as relative fluorescence units (RFU). (C) After six days of differentiation, macrophages were stimulated with heat-inactivated *S. aureus* and, after 24 hours of incubation, the supernatant was placed in the bottom chamber of a transwell system. HUVEC cells were placed in the top chamber and the system was incubated for 24 hours. HUVEC migrated cells were stained with DAPI, imaged and counted. (D) Histograms of the HUVEC migration quantification. (E) After six days of differentiation, in order to measure ROS generation, macrophages were stimulated with PMA (500 ng/mL) and luminol was added immediately prior to acquisition and reading. ROS production was monitored by chemiluminescence for 2 hours and expressed as relative light units (RLU). Graphs show mean ± SEM corresponding to up to three independent experiments. (F) Histogram representation of the Area Under the Curve (AUC) of the data shown in E. Graphs show the mean ± SEM of at least three independent experiments. For A, *n*=11 #*p* < 0.05, ##*p* < 0.01, ###*p*<0.001. Wilcoxon matched-pairs signed-rank test (#: GM-CSF vs. M-CSF-differentiated cells). For B, *n* = 8; for D, *n* = 8; for F, *n* = 7; #,\**p* < 0.05, ##,\*\**p* < 0.01. Wilcoxon matched-pairs signed-rank test (#: GM-CSF vs. M-CSF-differentiated cells; *: cells stimulated with β-glucan vs. non stimulated with β-glucan).

We therefore analysed and compared β-glucan-imprinted M-CSF- and GM-CSF-macrophages with regards to other specific functions that are hallmarks of the types of macrophages. M-CSF-M2-like macrophages have higher phagocytic affinities and capacities than M1-like macrophages (20), a process that we observed in our inactivated-bacteria phagocytosis assay (**Figure 3B**). Interestingly, β-glucan imprinting significantly reduced the quantity of phagocytosed bacteria by M-CSF-macrophages (**Figure 3B**). More specifically, less M-CSF-macrophages phagocytosed bacteria upon β-glucan reprogramming (**Supplementary Figure 3A**) and β-glucan-treated-M-CSF-macrophages engulfed less bacteria than their naïve counterparts (**Supplementary Figure 3B**). In fact, resulting β-glucan-imprinted macrophages showed the same phagocytic capacities as GM-CSF-macrophages.

M-CSF-macrophages are inducers of endothelial migration that is essential to angiogenesis, mainly through the production of VEGF, the most active endogenous pro-angiogenic factor in humans (21, 22). Given the observed modulation of VEGF by β-glucan imprinting (**Figure 1C**), we monitored the angiogenesis properties of our macrophages using conditioned media on HUVEC cells. As expected, we observed higher migration of HUVEC cells when cultured with conditioned media from M-CSF-macrophages compared to that of GM-CSF-macrophages (**Figure 3C, D**). β-glucan imprinting, which decreases VEGF expression in M-CSF-macrophages (**Figure 1G**) also decreased the migration-inducing properties of the conditioned media. Interestingly, β-glucan imprinting increased the migration properties of GM-CSF-macrophages conditioned media (**Figure 3C, D**) without increasing VEGF expression (**Supplementary Figure 4**). Ultimately supernatants from both types of β-glucan-imprinted-macrophages showed similar angiogenesis properties (**Figure 3D**).

ROS production is a hallmark of macrophage-mediated inflammatory response – ranging from antimicrobial activity, redox regulation of immune signalling to induction of inflammasome activation (23). As expected, ROS production was higher in GM-CSF-macrophages than M-CSF-macrophages (**Figure 3E, F**). β-glucan imprinting increased the capacity of M-CSF-macrophages to produce ROS to a similar level to that of GM-CSF-macrophages.

Altogether these results demonstrate that β-glucan imprinting differentially affects M-CSF-macrophages and GM-CSF-macrophages. The differentially decrease or increase of specific functions lead towards macrophages with similar phagocytosis, angiogenesis and ROS signalling properties.

### β-glucan reprogramming affects the expression of the master regulators of macrophage polarisation IRF5 and IRF3

To determine how this impediment of macrophage terminal differentiation occurs, we investigated the mechanism of action of β-glucan on the commitment of M-CSF- and GM-CSF-macrophages. β-glucan-induced innate immune memory acts on human primary monocytes (1) and macrophages (9) through the C-type lectin receptor Dectin-1. As described above, β-glucan imprinting reduces the TNFα responsiveness of GM-CSF macrophages whereas it exacerbates the TNFα responsiveness of M-CSF-macrophages (**Figure 4A**). Blocking the Dectin-1 receptor with an anti-Dectin-1 antibody alleviated these opposite effects of β-glucan on GM-CSF- and M-CSF-macrophages (**Figure 4A**). With respect to anti-inflammatory cytokines, as observed above, β-glucan imprinting only affects M-CSF-macrophages by reducing their expression of IL-10 (**Figure 4A**). Blocking the Dectin-1 receptor alleviated the β-glucan imprinting effect on M-CSF macrophages which turned out to secrete similar IL-10 cytokine levels upon stimulation as their naïve control counterparts (**Figure 4A**). These results demonstrate that β-glucan imprinting effects on polarising macrophages are mediated by Dectin-1 as already reported in human primary monocytes and macrophages (1, 9).

**Figure 4.**
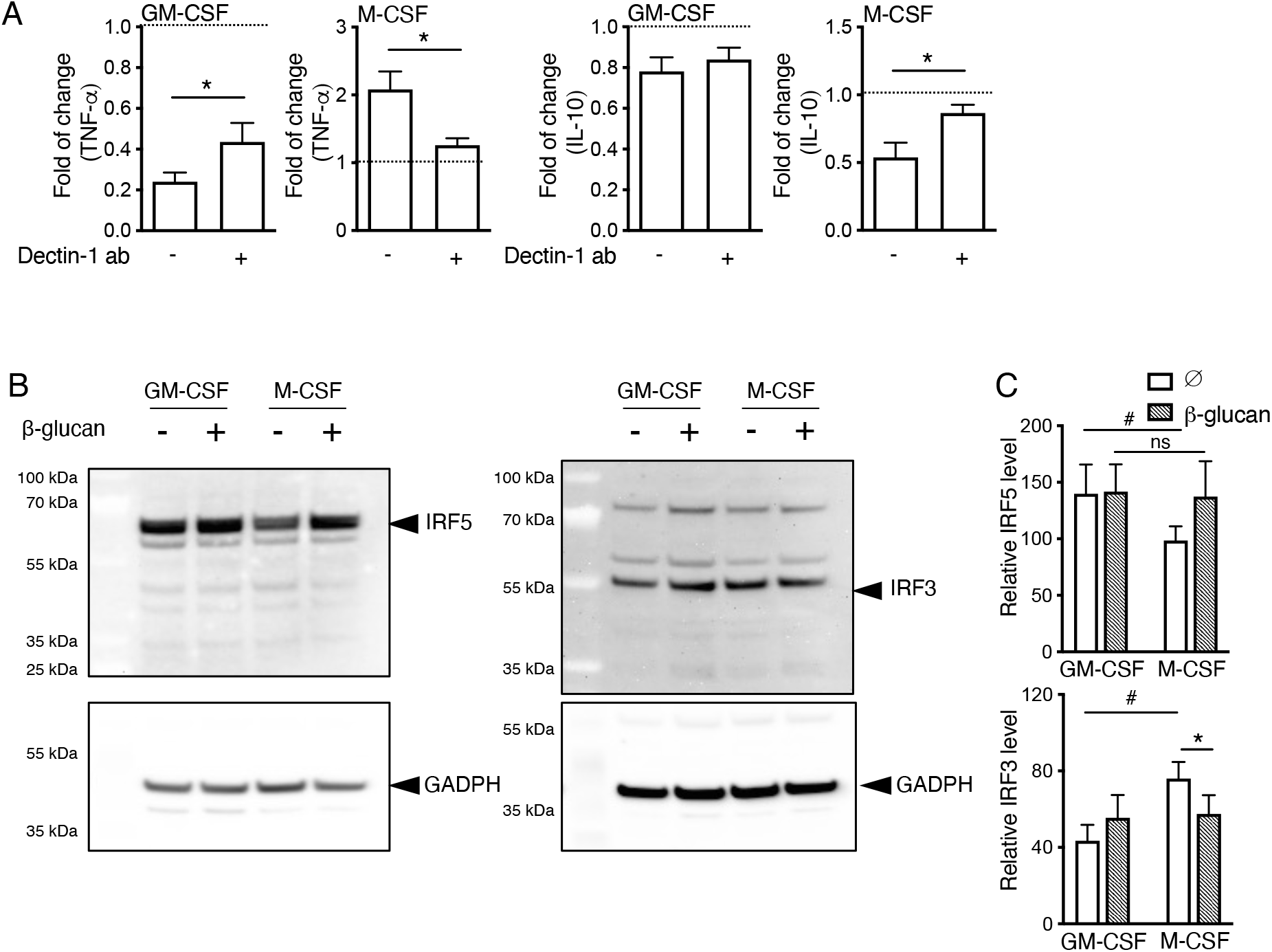
β-glucan imprinting is mediated by Dectin-1 and modulate the expression of IRFs. (A) Monocytes were pretreated for 30 minutes with an anti–Dectin-1 mAb or an IgG isotype control, then they were incubated with β-glucan or left untreated in a medium containing either GM-CSF or M-CSF. After 24 hours, the stimulus was washed away, and the cells were differentiated for an additional 5 days, after which the macrophages were restimulated with heat-inactivated *S. aureus*. Culture supernatants were collected following 24h restimulation and the concentrations of secreted TNF-α and IL-10 were determined by ELISA. The results obtained for the cells differentiated in the presence of β-glucan were normalised by its respective control; results above one indicate an increase mediated by β-glucan compared to control; results below one indicate a decrease mediated by β-glucan compared to control. (B) Monocytes were differentiated with GM-CSF or M-CSF and left untreated or stimulated with β-glucan for the first 24 hours. After this first period, the stimulus was washed out and cells were differentiated for another five days with GM-CSF or M-CSF, with the medium being refreshed once during this period. At day 6 of the differentiation process, macrophages were re-stimulated with *S. aureus* for 24 hours. Total cellular protein content was obtained and IRF5 and IRF3 expression was assessed by western blot. GAPDH was used as loading control. Blots are representative of five independent experiments. (C) Densitometric analysis of the blots shown in (B). Data represent mean values ± SEM of the analysis of at least three independent experiments. For A, *n* = 5; for C, *n* = 7; *,#*p* < 0.05. Wilcoxon matched-pairs signed-rank test. (#: GM-CSF vs. M-CSF-differentiated cells; *: cells stimulated with β-glucan vs. non stimulated with β-glucan).

Members of the interferon-regulatory factor (IRF) family determine commitment to GM-CSF-M1-like or M-CSF-M2-like macrophages. The transcription factor IRF5 is the master regulator of M1 macrophage polarisation (24). As expected, IRF5 was more expressed in human GM-CSF-M1-like-macrophages compared to M-CSF-M2-like macrophages (**Figure 4B, C)** (24). Of interest, β-glucan imprinting of M-CSF-macrophages increased the expression of the GM-CSF-master regulator IRF5 to the same level as in GM-CSF macrophages. The transcription factor IRF3 is predominantly activated in M-CSF-M2-like macrophages (25) and is associated with anti-inflammatory microenvironments and contributes to the polarisation towards a M2 macrophage phenotype (26). As expected, IRF3 was found to be more expressed in our M-CSF-macrophages compared to GM-CSF-cells (**Figure 4B,C**). Interestingly again, β-glucan imprinting significantly decreased the expression of IRF3 in M-CSF-macrophages, to the same level as in GM-CSF macrophages. Altogether, β-glucan reprogramming affected the specificity of macrophage polarisation through Dectin-1 and the modulation of the expression of the master regulators of the macrophage polarisation IRFs.

### β-glucan imprinting skews the commitment of macrophages hastily during the differentiation process

We observed that β-glucan imprinting of macrophages in the first 24 hours affects their differentiated features 6 days after, which could be seen as either skewing or reverting polarisation. To test these hypotheses, we next assessed whether β-glucan imprinting rapidly skews the commitment of macrophages, reprogramming them from the onset towards imprinting, or whether it progressively reverts the polarisation over time. We measured the cytokine responses of the cells to heat-inactivated *S. aureus* stimulation as a hallmark of differentiation. Measurements were performed as usual, 6 days after β-glucan treatment, but also before β-glucan imprinting (monocytes), a day after, and 3 days after (**Figure 5A**).

**Figure 5.**
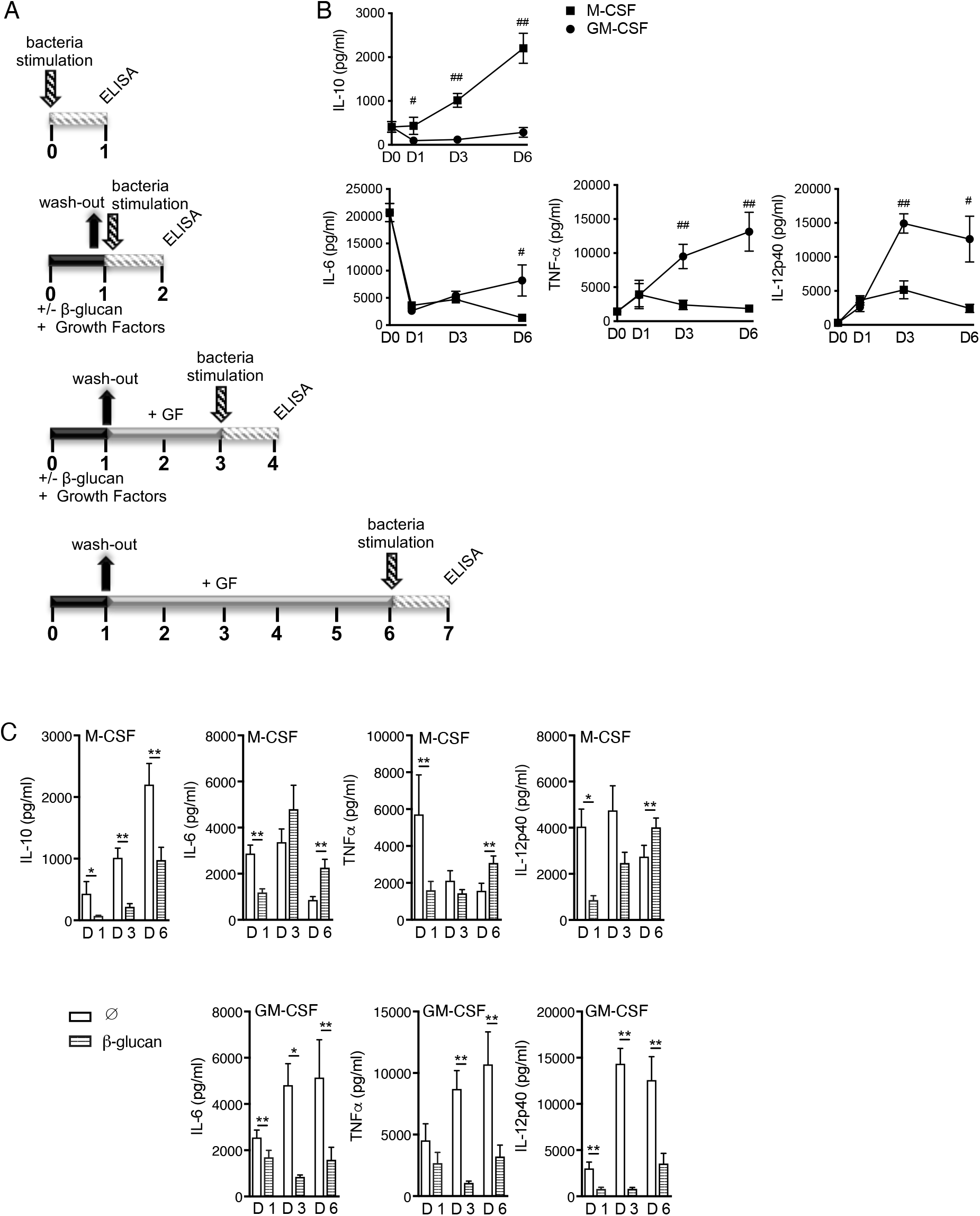
β-glucan imprinting modulates the release of cytokines already during the differentiation process. (A) Schematic overview of the experiment. Four different timelines of differentiation and stimulation were designed to evaluate the response of the cells during the process. Culture supernatants were always collected after 24h of *S. aureus* stimulation. (i) Day 0: Monocytes were isolated and stimulated the same day with *S. aureus*. (ii) Day 1: Monocytes were differentiated with GM-CSF or M-CSF, in the absence or presence β-glucan for 24 hours. After this period, the system was washed before stimulation with *S. aureus*. (iii) Day 3: Monocytes were differentiated with GM-CSF or M-CSF, in the absence or presence β-glucan for 24 hours. After the first 24 hours, the system was washed away, and cells maintained in GM-CSF or M-CSF. On day 3 of the differentiation process the cells were stimulated with *S. aureus*. (iv) Day 6: Monocytes were differentiated with GM-CSF or M-CSF, in the absence or presence β-glucan for 24 hours. After the first 24-hour period, the system was washed away, and cells maintained in GM-CSF or M-CSF medium. On day 3 GM-CSF and M-CSF medium was refreshed. At day 6, the macrophages were stimulated with *S. aureus*. GF=growth factors. (B,C) Culture supernatants were collected after 24h of *S. aureus* stimulation at the different time-points depicted in (A) and concentrations of secreted TNF-α, IL-6, IL-12, and IL-10 were determined by ELISA. Graphs show the mean ± SEM of at least three independent experiments; *n* = 8, #,\**p* < 0.05, **,##*p* < 0.01. Wilcoxon matched-pairs signed-rank test. (#: GM-CSF vs. M-CSF-differentiated cells; *: β-glucan-imprinted vs. non stimulated with β-glucan).

First, without β-glucan input, it is interesting to note that with respect to the pro-inflammatory cytokines, none of the differentiating macrophages were polarised (i.e. GM-CSF > M-CSF) after only one day of incubation with growth factors (**Figure 5B)**. Polarisation (GM-CSF > M-CSF) started to be observable for the pro-inflammatory IL-12 and TNFα from day 3 and was maintained throughout the differentiation, up to day 6. For IL-6, although differentiating macrophages already expressed lower cytokines than monocytes from day 1, the polarisation required more time and was only observable at day 6. So altogether polarisation for pro-inflammatory cytokines is a progressive event that occurs over time. In contrast to pro-inflammatory cytokines, polarisation with respect to the anti-inflammatory cytokine IL-10 (M-CSF > GM-CSF) was observable from day 1 and maintained throughout the experiment (**Figure 5B**).

The decrease in the anti-inflammatory IL-10 cytokine detected in M-CSF-macrophages upon β-glucan imprinting was observable at the onset from day 1 (**Figure 5C**) and was maintained over time. With respect to pro-inflammatory signature, β-glucan imprinting affected both types of macrophages early and inhibited their responses to *S. aureus* (**Figure 5C**). This inhibition at the onset was maintained for GM-CSF over the time of the experiment, culminating in the decreased responsiveness described above at day 6 (**Figure 5C**). However, for M-CSF, the final reprogramming effect with increased responsiveness only appeared after 6 days post-treatment (**Figure 5C**), only after control naïve M-CSF-macrophages reached their final polarisation features. Altogether, these results highlight that β-glucan affects the polarisation and differentiation processes of GM-CSF- and M-CSF-macrophages as soon as it has been detected by the cells, skewing their full differentiation in either direction. β-glucan-imprinted-macrophages are therefore less terminally differentiated, but not because of a reversion and rather because of an impediment of the primary processes of polarisation.

## Discussion

The imprinting of human primary monocytes associated with enhanced pro-inflammatory responsiveness towards several different pathogens suggest a potential therapeutic role for β-glucan-induced innate immune memory in diverse infectious disease context. However, during infection, bloodstream classical monocytes are mostly circulating intermediates. In tissues, where they are recruited upon infections, monocytes differentiate or polarise into macrophages following conditioning by local host factors, and microbial products. Therefore, the interaction of β-glucan with innate cells *in vivo* would occur in an evolving environment. Gaining insights on the modulation properties of β-glucan imprinting in complex physiological environments is important for envisioning future therapeutics harnessing the beneficial effects of innate immune memory. We therefore proposed in this study to investigate the specific effects of β-glucan imprinting on polarising macrophages in complex environment.

Consistent with previous studies investigating *in vitro* human monocytes imprinting by β-glucan with human serum containing low but detectable amount of M-CSF, we observed that β-glucan imprinting of polarising macrophages with higher doses of exogenous M-CSF increased their pro-inflammatory cytokines response. However, and never reported before, we observed that β-glucan imprinting of GM-CSF-differentiating macrophages decreased their responsiveness, resulting in less inflammatory macrophages. In addition, although the M-CSF environment that we used is the most resembling milieu in the bloodstream, we still observed specific effects. As such, M-CSF-macrophages imprinted with β-glucan present a lower phagocytic index that is in contrast to human serum-cultured monocytes (1). These observations highlight the importance of the environment surrounding the innate cells outside of the bloodstream on the effects of β-glucan imprinting and enlarge the prospects of using innate immune memory in future clinic applications

We consistently observed that β-glucan reprogramming differentially imprints GM-CSF- and M-CSF-macrophages towards a similar type of macrophage that presents both intermediate pro- and anti-inflammatory responses. β-glucan reprogramming confers a unique phenotype to differentiating macrophages in which the specific markers of terminally differentiated macrophages we measured were decreased in the related macrophages. These observations strongly suggest that β-glucan reprogramming of differentiating macrophages provide cells with larger panel of reactivity to secondary input, potentially making them more plastic in terms of the type of response required. This could explain the previously reported aspecifity of the response of β-glucan-imprinted cells towards different non-related pathogens (1).

The decrease differentiation marks in β-glucan imprinted and potentially associated increased plasticity suggest that β-glucan imprinting of macrophages in complex environment could affect non-infectious related host-mechanisms. Indeed, macrophages functions are not dedicated to immunity against pathogens. We observed that β-glucan imprinting, which decreases VEGF expression in M-CSF-macrophages also decreased the HUVEC migration-inducing of the conditioned media. Excessive VEGF and subsequent angiogenesis have been associated with atherosclerotic plaque progression (27, 28) and tumour angiogenesis (29). β-glucan imprinting might therefore decrease risk of cardiovascular disease or tumour progression. Of interest, β-glucan imprinting increased the migration properties of GM-CSF-macrophages conditioned media without increasing VEGF expression. Angiogenesis plays critical roles in human physiology that range from reproduction and foetal growth to wound healing and tissue repair (30). Hence β-glucan imprinting could represent therapeutic opportunities to potentiate the beneficial functions of angiogenesis induced by GM-CSF-macrophages secretome, independently of VEGF.

β-glucan imprinting of M-CSF- and G-MCSF-macrophages is mediated by the Dectin-1 receptor as demonstrated by our use of the anti-Dectin-1 antibody. Of note, Dectin-1 expression is highly up-regulated by GM-CSF (31). This increased amount of Dectin-1 receptor at the surface of GM-CSF-macrophages might not be fully blocked by the anti-Dectin-1 antibody, compared to M-CSF-macrophages, which could explain why the restoration of TNFα secretion in Dectin-1-inhibited-GM-CSF-macrophages is not total. Overall, these results together with previous studies assert Dectin-1 as a master regulator of β-glucan imprinting in human primary mononuclear phagocytes.

Based on the CD4 T helper type 1 (Th1), Th2 and Th17 cells polarisation paradigm (32), M1-GM-CSF macrophages produce proinflammatory cytokines, mediate resistance to pathogens and drive a Th1/Th17 response, whereas M2-M-CSF macrophages produce anti-inflammatory cytokines, promote tissue repair and remodelling and drive a Th2 response. In M1 macrophages, IRF5 dictates the expression of proinflammatory genes such as *IL-12* whilst repressing anti-inflammatory genes like *IL-10,* driving a potent Th1/Th17 response (24). Here, β-glucan reprogramming of M-CSF-M2-like-macrophages increased IRF5 expression, with a concomitant increase in IL-12 expression and decrease in IL-10 expression, therefore driving the β-glucan**-**M-CSF-macrophages towards setting up an environment for a CD4 Th1/ Th17 response. This suggests that not only does β-glucan imprinting skew macrophages differentiation but could also shape the subsequent Th responses. This potentially add another layer of plasticity triggered by β-glucan reprogramming. Indeed, there is evidence that all Th cells, with the exception perhaps of Treg cells, retain a certain degree of plasticity upon differentiation into effector cells (33). Although prolonged stimulation induces a more stabilized Th1 or Th2 program, polarized Th cells retain flexibility with regard to their transcriptional signature for several rounds of expansion, giving them enough time to adjust their response to stimulation. β-glucan reprogramming, which is providing such plasticity to macrophages, might in turn influence the Th program and plasticity.

In conclusion, we have established that complex environment strongly impacts the effects of β-glucan reprogramming of phagocytes. Not only does β-glucan imprinting enhance responsiveness and function of macrophages but under certain conditions it can reduce it. In addition, β-glucan imprinting skew the polarisation macrophages leading towards a common type of less differentiated macrophages. β-glucan-induced innate memory in macrophages imprints their terminal differentiation through the Dectin-1 receptor and a concomitant alteration of master regulators IRF5 and IRF3 expression. Deciphering how β-glucan specifically imprints polarising macrophages in complex environments provides fundamental knowledge bringing us a step further towards exploiting its modulatory properties.

## Acknowledgments

The authors wish to thank Dr David L Williams for generously providing the purified β-glucan.

## Disclosures

The authors have no financial conflicts of interest.

## Supplementary Materials for

**Supplementary Figure 1.**
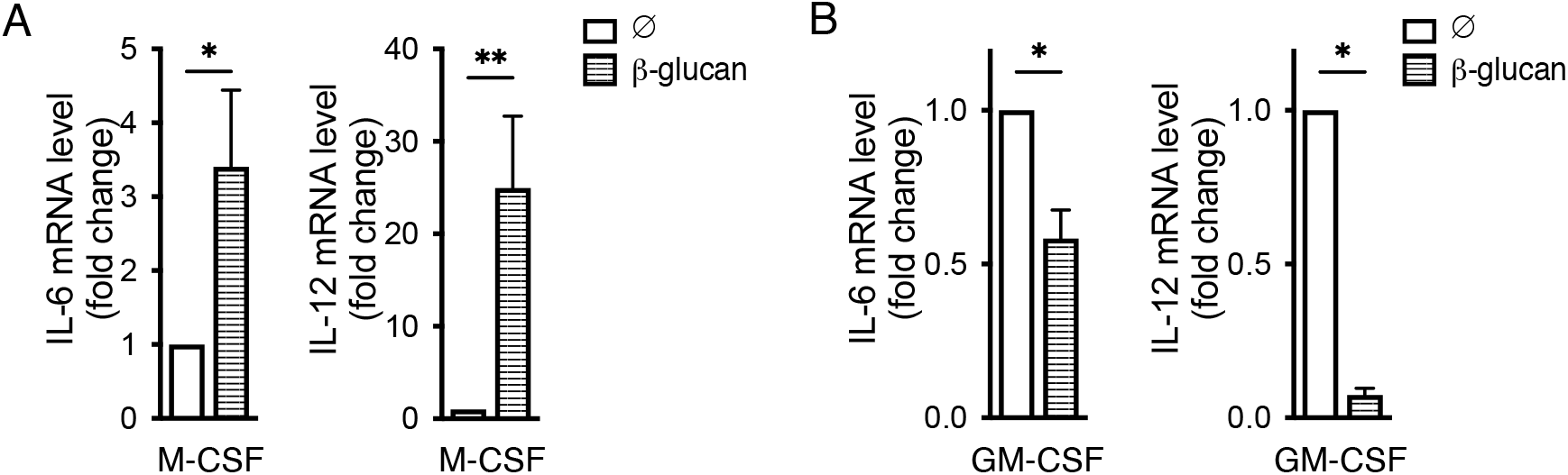
Transcriptional regulation of the β-glucan imprinting on differentiating macrophages. Monocytes were differentiated with M-CSF (A) or GM-CSF (B) in the absence or presence of β-glucan for the first 24 hours. After the first 24 hours, cells were maintained in medium enriched in the related growth factor that was changed once during this second period. At day 6 of differentiation, the macrophages were re-stimulated with heat-inactivated *E. coli*. After 3 hours of stimulation, RNA was extracted from macrophages and the expression of *IL6* and *IL12* evaluated by RT-qPCR. Graphs show the mean ± SEM of at least three independent experiments; *n* = 7; \**p* < 0.05, \*\**p* < 0.01. Wilcoxon matched-pairs signed-rank test.

**Supplementary Figure 2.**
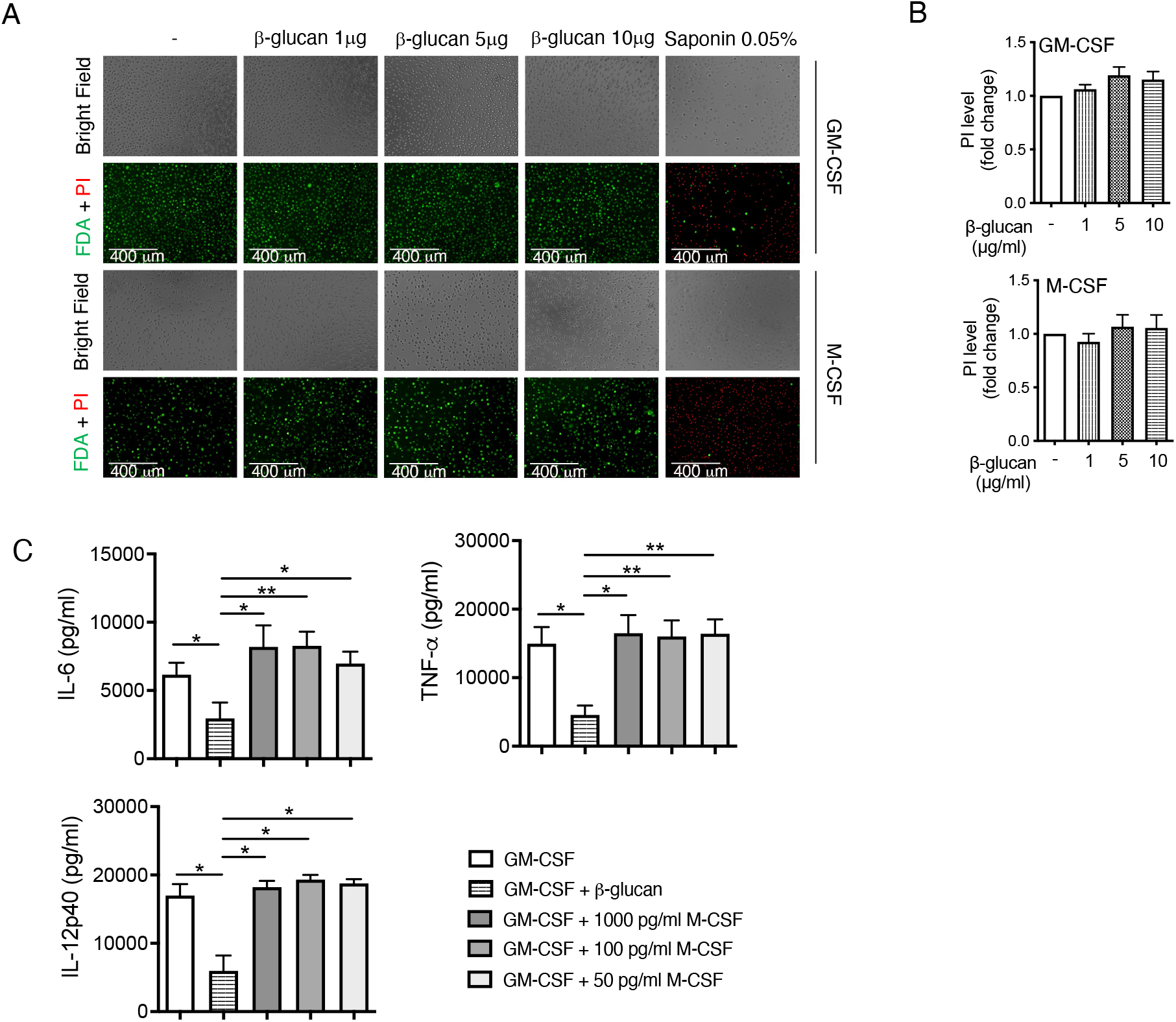
β-glucan imprinting is not mediated by cell death or increased M-CSF secretion. (A) Monocytes were preincubated in the presence or absence of β-glucan and then differentiated with either GM-CSF or M-CSF. At day 6 of differentiation, live/dead rate was measured in the resulting macrophages by loading cells with Fluorescein diacetate (FDA) (green, live cells) and Propidium Iodide (PI) (red, dead cells). Saponin 0.05% was used as a positive control for the PI staining. (B) Quantification of PI level relative to the control. (C) Monocytes were preincubated with β-glucan or the indicated concentration of M-CSF in a medium containing GM-CSF. After 24 hours, stimuli were washed away, and the cells were differentiated for an additional 5 days in the presence of GM-CSF. At day 6 of differentiation, resulting macrophages were restimulated for 24h with heat-inactivated *S. aureus,* culture supernatants were collected, and the concentrations of secreted IL-6, TNF-α, and IL-12p40 were determined by ELISA. Graphs show the mean ± SEM of at least two independent experiments. For B, *n* = 4; for C, *n* = 3 \**p* < 0.05, \*\**p* < 0.01. Wilcoxon matched-pairs signed-rank test.

**Supplementary Figure 3.**
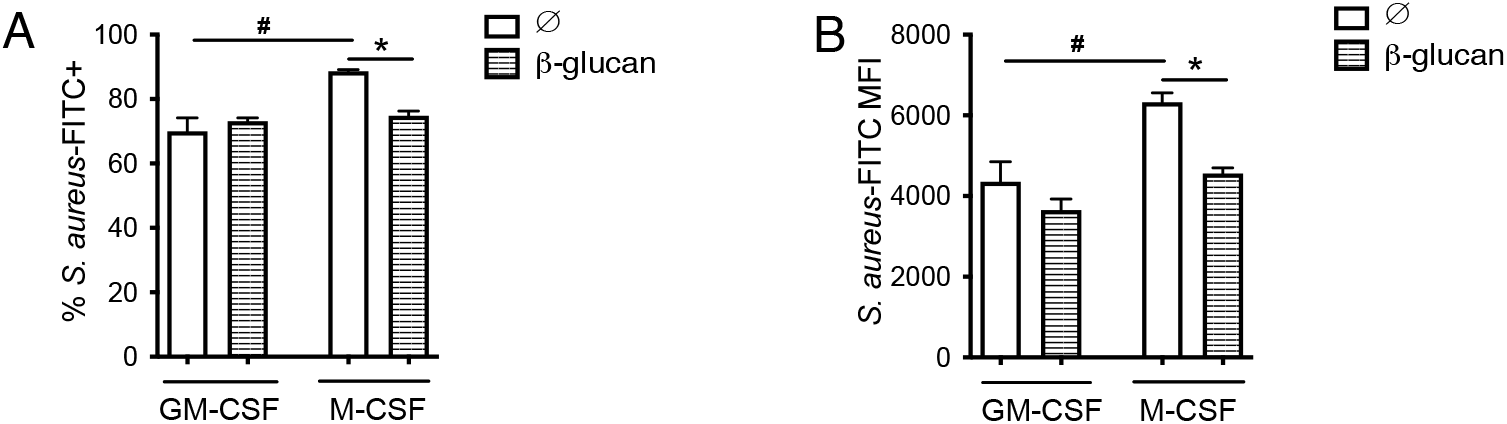
Phagocytosis rate and efficacy are both modulated by β-glucan-induced innate memory on polarising macrophages. Monocytes were preincubated in the presence or absence of β-glucan and then differentiated in macrophages with either GM-CSF or M-CSF. After 6 days of culture, resulting macrophages were re-stimulated with pHrodo™ Green *S. aureus* Bioparticles™ for 30 min. Cells were then detached and analysed by flow cytometry to determine (A) the percentage of stained cells and (B) the mean fluorescence intensity (MFI). Graphs show mean ± SEM; *n* = 3 independent experiments. ^#,^\**p* < 0.05. Wilcoxon matched-pairs signed-rank test (#: GM-CSF vs. M-CSF-differentiated cells; *: cells stimulated with β-glucan vs. non stimulated with β-glucan).

**Supplementary Figure 4.**
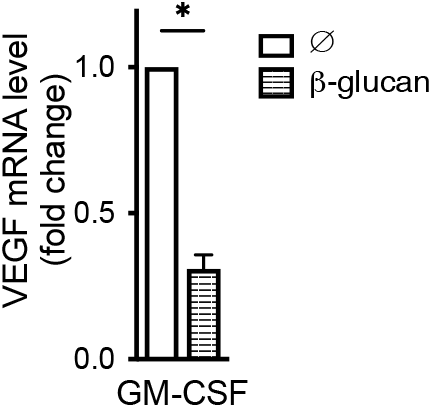
β-glucan-imprinting of GM-CSF-polarising macrophages do not increase ***VEGF*** expression. Monocytes were preincubated in the presence or absence of β-glucan and then differentiated in macrophages with GM-CSF. At day 6 of differentiation, macrophages were re-stimulated for 18h with heat-inactivated *S. aureus* and *VEGF* gene expression was analysed by Real-Time quantitative PCR (RT-qPCR). Results were normalised to β-actin expression levels. The results obtained for the cells differentiated with GM-CSF in the absence of β-glucan were arbitrarily set at 1 to express the results as fold change. Graphs show mean ± SEM; *n* = 3 independent experiments; *n* = 7. \**p* < 0.05. Wilcoxon matched-pairs signed-rank test.

## Notes

1 A.C.P, G.C., M.B., R.L., and J.Q. were supported by the ANR JCJC grant (ANR-16-CE15-0014-01), and by Institut Carnot–Microbes et Santé grant (n°11 CARN-017-01) (to J.Q.). The authors also acknowledge funding from CAPES finance code 001. A.C.P. was supported by the CAPES-COFECUB Franco-Brazilian Research Exchange Program (88887.281641/2018-00). For the purpose of open access, the author has applied a CC BY public copyright license to any author accepted manuscript version arising from this submission.

### Competing Interest Statement

The authors have declared no competing interest.

### Summary of Updates

This revision is a full-length version of the article.

